# Structure, interaction, and nervous connectivity of beta cell primary cilia

**DOI:** 10.1101/2023.12.01.568979

**Authors:** Andreas Müller, Nikolai Klena, Song Pang, Leticia Elizabeth Galicia Garcia, Oleksandra Topcheva, Solange Aurrecoechea Duran, Davud Sulaymankhil, Monika Seliskar, Hassan Mziaut, Eyke Schöniger, Daniela Friedland, Nicole Kipke, Susanne Kretschmar, Carla Münster, Jürgen Weitz, Marius Distler, Thomas Kurth, Deborah Schmidt, Harald F. Hess, C. Shan Xu, Gaia Pigino, Michele Solimena

## Abstract

Primary cilia are sensory organelles present in many cell types. Based on an array of microtubules termed axoneme they form a specialized membrane compartment partaking in various signaling processes. Primary cilia of pancreatic islet beta cells play a role in autocrine and paracrine signaling and are linked to diabetes. Yet, the structural basis for their functions is unclear. We present three-dimensional reconstructions of complete mouse and human beta cell cilia, revealing a disorganized 9+0 axoneme structure. Within the islet, cilia are spatially confined within deep ciliary pockets or squeezed into narrow extracellular spaces between adjacent cells. Beta and alpha cell cilia physically interact with neighboring islet cells pushing and strongly bending their plasma membranes. Furthermore, beta cells can contain multiple cilia that can meet with other islet cell cilia in the extracellular space. Additionally, beta cell cilia establish connections with islet-projecting nerves. These findings highlight the pivotal role of beta cell primary cilia in islet cell connectivity, pointing at their potential functional role in integrating islet intrinsic and extrinsic signals. These novel insights contribute to understanding their significance in health and diabetes.

## Introduction

Pancreatic islet beta cells control blood glucose levels by secreting the hormone insulin. Insulin secretion is a concerted process regulated by autocrine signaling cascades, paracrine interactions with other beta and non-beta pancreatic islet cells as well as factors of systemic and neuronal origin (1; 2; 3). The specific architecture of the islets plays a significant role in the orchestration of these pathways, as these mini-organs are highly vascularized (4), with their endocrine cells forming rosettes around the capillaries (5). Furthermore, islets make multiple connections with the nervous system (3). Beta cells have a special morphology with three distinct domains: a vascular pole, lateral poles and an a-vascular pole, which usually is defined by the projection of a single primary cilium (6). A deeper understanding of this highly specialized compartment in beta cells is especially desirable in view of its proposed involvement in the pathogenesis of diabetes - a common metabolic disorder resulting from impaired beta cell insulin secretion.

The majority of cell types in our body have one primary cilium which can act as a “cellular antenna” for the reception of paracrine signals, and cilia dysfunction has been linked to a number of diseases usually referred to as ciliopathies such as Bardet-Biedl-syndrome or polycystic kidney/liver disease (7; 8). Cilia are microtubule-based organelles that extend from the cell forming a specialized membrane compartment (9). The underlying microtubule array - the axoneme - extends from the basal body, which is constructed out of nine microtubule triplets (A-, B- and C-tubule). Furthermore, in many cell types the base of the ciliary membrane forms a distinct domain termed ciliary pocket (10; 11). Cilia can be roughly divided into motile cilia (present in airways, sperm tails etc.) and non-motile cilia, referred to as primary cilia (9). Motile cilia have a complex and stereotyped structure with 9 microtubule doublets (A- and B-tubule) organized in a 9-fold symmetry at the periphery of the axoneme and 2 singlet microtubules at the center (9+2) (12). This microtubular structure is the frame to which all the motility protein complexes, such as axonemal dyneins and radial spokes, bind to and work on to generate the ciliary beating (13; 14; 15; 16; 17). Diversely, the microtubule structure of primary cilia typically follows a 9+0 scheme with a ring of 9 microtubule doublets (18). However, serial section electron microscopy (EM) data have suggested the breaking of the 9 fold symmetry in close proximity to the basal body (19; 20) leading to a variable axoneme structure (21). Recently, reconstructions of complete primary cilia from cultured kidney cells revealed substantial variations from the 9+0 architecture, with the progressive loss of microtubule doublets, as well as the B-tubule termination as the axoneme progresses distally (22; 23). Furthermore, axonemal microtubules consistently display a distortion from a radial arrangement, to a more bundled organization, with one or more singlets or doublets moving away from the axoneme and to the center of the lumen. These data indicate that the 9 fold symmetry is not as relevant for the signaling function of primary cilia, as it is for the motility function in motile cilia.

Primary cilia of beta cells have been detected by various microscopy methods (24; 25; 26; 27; 28; 29; 30). They contribute to paracrine signal transduction within the islets of Langerhans (31). Furthermore, beta cell cilia play a pivotal role in glucose homeostasis and their loss or dysfunction impairs insulin secretion and leads to abnormal beta cell polarity (32). Perturbation of the expression of ciliary genes has been linked to the development of diabetes mellitus (33; 34). Moreover, in a mouse model of type 2 diabetes ciliogenesis appears to be impaired (34). Recently, it has been postulated that beta cell primary cilia can actively move in response to glucose stimulation (35). However, the structural basis for this phenomenon is currently unclear. Three-dimensional (3D) reconstructions of the axoneme of beta cell primary cilia are currently not available. Until now the ultrastructure of beta cell primary cilia and their axonemes has been investigated only by two-dimensional (2D) transmission electron microscopy (TEM) (36; 35; 24; 26) as well as scanning electron microscopy (SEM) (30). Some of these studies indicate a dis-organization of the axoneme. However, the complete arrangement of the microtubules of the primary cilium cannot be extrapolated from 2D images. Ultimately, full 3D reconstructions of beta cell primary cilia are necessary to understand their role in islet signaling. In addition to gathering information about the structure of the axoneme, direct visualization of primary cilia in the islet is of direct interest to gain insight into ciliary involvement in islet function.

In this study we have employed two volume electron microscopy (vEM) methods with 3D segmentation accompanied by ultrastructural expansion microscopy (U-ExM) to investigate the 3D structure of primary cilia of mouse and human beta cells in the context of isolated islets and pancreas tissue. We found that the axoneme of beta cell cilia in mice and humans is disorganized. We furthermore observed that beta cell primary cilia are spatially confined either within long ciliary pockets or restricted by surrounding cells, including acinar cells and islet vasculature. We also discovered close interactions of primary cilia originating from beta or alpha cells with neighboring islet cells or their cilia. Finally, we found that islet cell primary cilia can connect to and even form synapses with the innervation of the pancreas. Our structural data enable a better understanding of the beta cell primary cilia as compartments for signal transduction within the islet, as well as with the exocrine tissue and the autonomic nervous system, contextualizing beta cell primary cilia as major connective hubs in islet function.

## Results

### The axoneme of mouse and human beta cell cilia follows a 9+0 organization which is not maintained over the whole cilia length

In order to resolve the organization of beta cell primary cilia it is pivotal to obtain high resolution 3D imaging data of their complete structure. This has so far been achieved for cells cultured in monolayers by serial section electron tomography of chemically fixed kidney cells (23; 22). Additionally, Kiesel et al. provided molecular data by cryo-electron tomography (cryo-ET) of kidney cilia identifying actin filaments within the axonemal structure and EB1 proteins decorating the lattice of the A-microtubules. However, high resolution 3D reconstructions of complete primary cilia and their axonemes within tissues are currently not available. This is due to their low abundance and slender morphology, which makes their targeting and localization in EM datasets a tedious task. To reconstruct mouse beta cell primary cilia in situ we reviewed our recently published data (28; 29; 37; 38; 39) as well as previously unpublished datasets obtained by focused ion beam scanning electron microscopy (FIB-SEM) of isolated mouse islets fixed by high pressure freezing (HPF) for the presence of primary cilia. We could indeed find primary cilia characterized by the presence of a basal body, axoneme and ciliary membrane (Fig. 1a, Supp.Fig. 1b). The voxel size of 4 nm was sufficient to discriminate microtubule doublets and singlets and the FIB-SEM volumes were large enough to contain complete primary cilia (Fig. 1a). This allowed for their 3D reconstruction (Fig. 1c, Video abstract) and assessment of their interaction with the neighboring tissue. Especially for this task, fixation with HPF enabled an optimal preservation of the ultrastructure because it avoids shrinking of the plasma membranes, as usually observed after chemical fixation. To obtain data on human beta cell primary cilia we processed chemically fixed pancreas specimens from pancreatectomized living donors (40; 41) and performed serial section electron tomography (ssET). Before the acquisition of electron tomograms, the serial sections were screened for the presence of basal bodies and axonemes by TEM (Supp. Fig. 1a). The corresponding regions were then imaged by ET and consecutive tomograms were aligned and stitched together. ET allows for imaging at higher resolution compared to FIB-SEM, so we decided for a voxel size of 1.307 nm in order to better resolve the microtubules as well as densities of other proteins of the cilia lumen. We could detect primary cilia in human beta cells and again clearly resolve microtubule doublets and singlets of the axoneme (Fig. 1b, Video 1). Out of vEM we mostly reconstructed the primary cilia of beta cells, which we identified by the morphology of their secretory granules (SGs), consisting of dense insulin cores surrounded by a translucent halo (42; 43; 44). We refer to these cilia as beta cell primary cilia, whereas cilia with unknown cell origin are referred to as islet cell primary cilia.

**Fig. 1.**
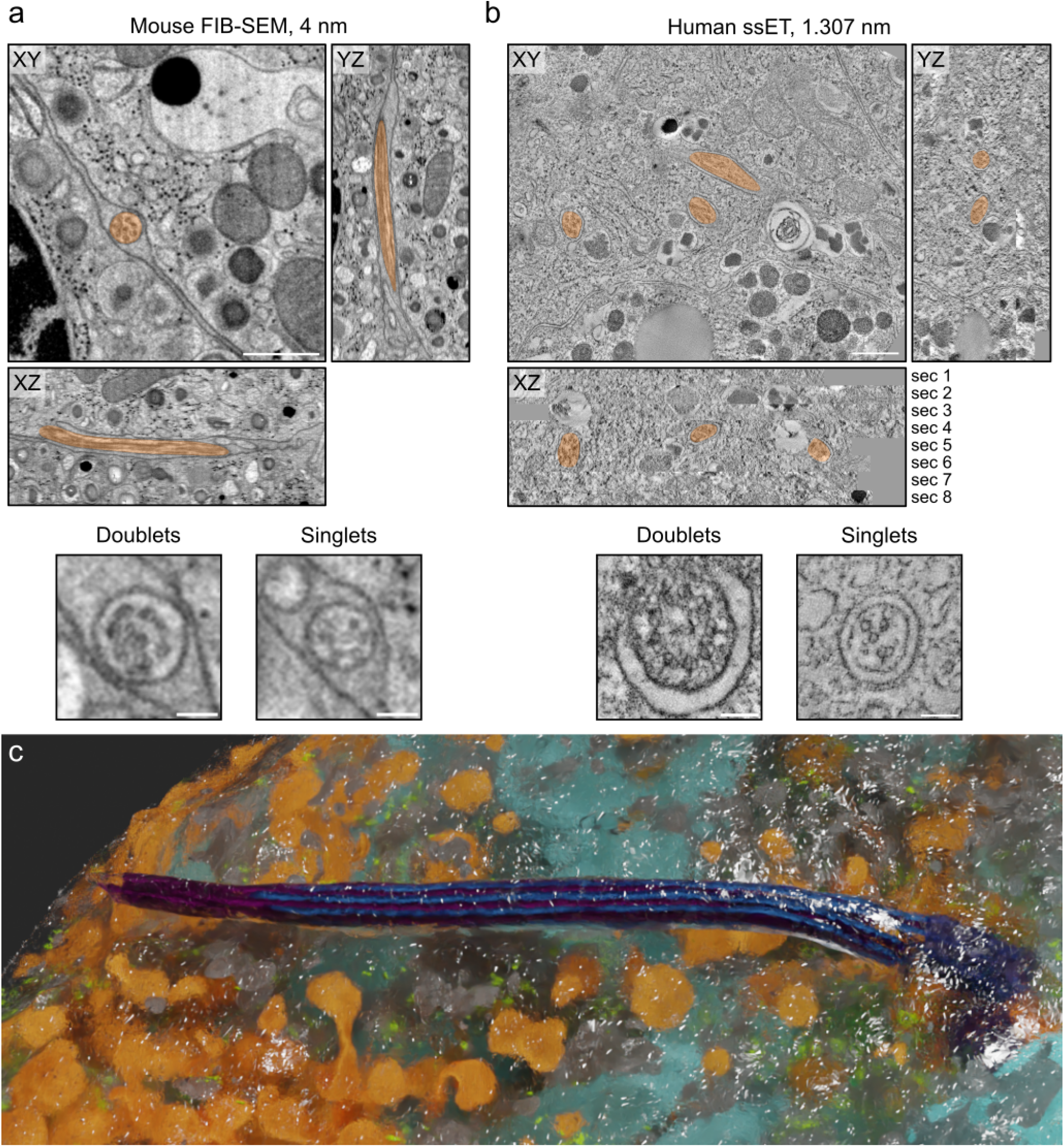
vEM resolves mouse and human beta cell primary cilia ultrastructure. FIB-SEM of isolated mouse islets **a)** and ssET comprising 8 consecutive sections of pancreas tissue of living human donors **b)**. Primary cilia are marked in orange. Both imaging modalities resolve the microtubule organization of the axoneme with microtubule doublets and singlets. Scale bars overview: 500 nm, scale bars axonemes: 100 nm. **c)** 3D rendering of a mouse beta cell primary cilium with microtubules (blue, purple), mitochondria (light blue), insulin SGs (orange), endoplasmic reticulum (gray), ribosomes (green), and plasma membrane (transparent).

Primary cilia in our vEM data showed a diverse morphology with straight as well as highly curved shapes (e.g. compare Fig. 2a, Fig. 4e). The length of primary cilia in vEM datasets of mouse islets and human pancreas was 4.2 μm and 2.7 μm, respectively. Segmentation of the microtubules of the axoneme of the vEM data and subsequent analysis revealed a strong deviation from the classical 9+0 organization, which was similar to that previously observed in 3D reconstructions of other cell types (23; 22) (Video 2). As expected, in both mouse and human beta cell primary cilia the C-tubules of the 9 triplets of the basal body terminated in the so-called transition zone whereas A- and B-tubules continued more distally as 9 doublets (Fig. 2a). Over the length of the cilium some of the doublets were displaced towards the center of the cilium breaking axoneme symmetry and making its organization appear even more dis-organized (Fig. 2b, c and d). During this transition one or two doublets passed the center of the cilium getting closer to the opposite side (Fig. 2c and d). The maximum distance of these doublets to the cilia membrane was approximately 100 nm whereas doublets that stayed close to the membrane had a distance of approximately 40 nm to it. In all reconstructed cilia the microtubule doublets were not present along the whole length of the cilia. Furthermore, we could not detect repetitive electron-dense structures compatible with the presence of motility components. vEM also enabled the precise measurement of the lengths of A- and B-tubules. In the mouse cilium A-tubules had a mean length of 4.57 ± 0.14 μm, and B-tubules a mean length of 3.03 ± 1.01 μm (Fig. 2e). In the fully reconstructed human beta cell primary cilium the length distribution was much more variable, with A-tubules measuring 2.05 ± 1.01 μm and B-tubules 1.20 ± 0.65 μm (Fig. 2e). In most cases the B-tubules terminated much before the A-tubules (Fig. 2a, c and d, Video 2), although we could observe cases in human beta cells where the opposite occurred (Supp. Fig. 2a). Furthermore, A- and B-tubules were slightly tortuous (Fig. 2f).

**Fig. 2.**
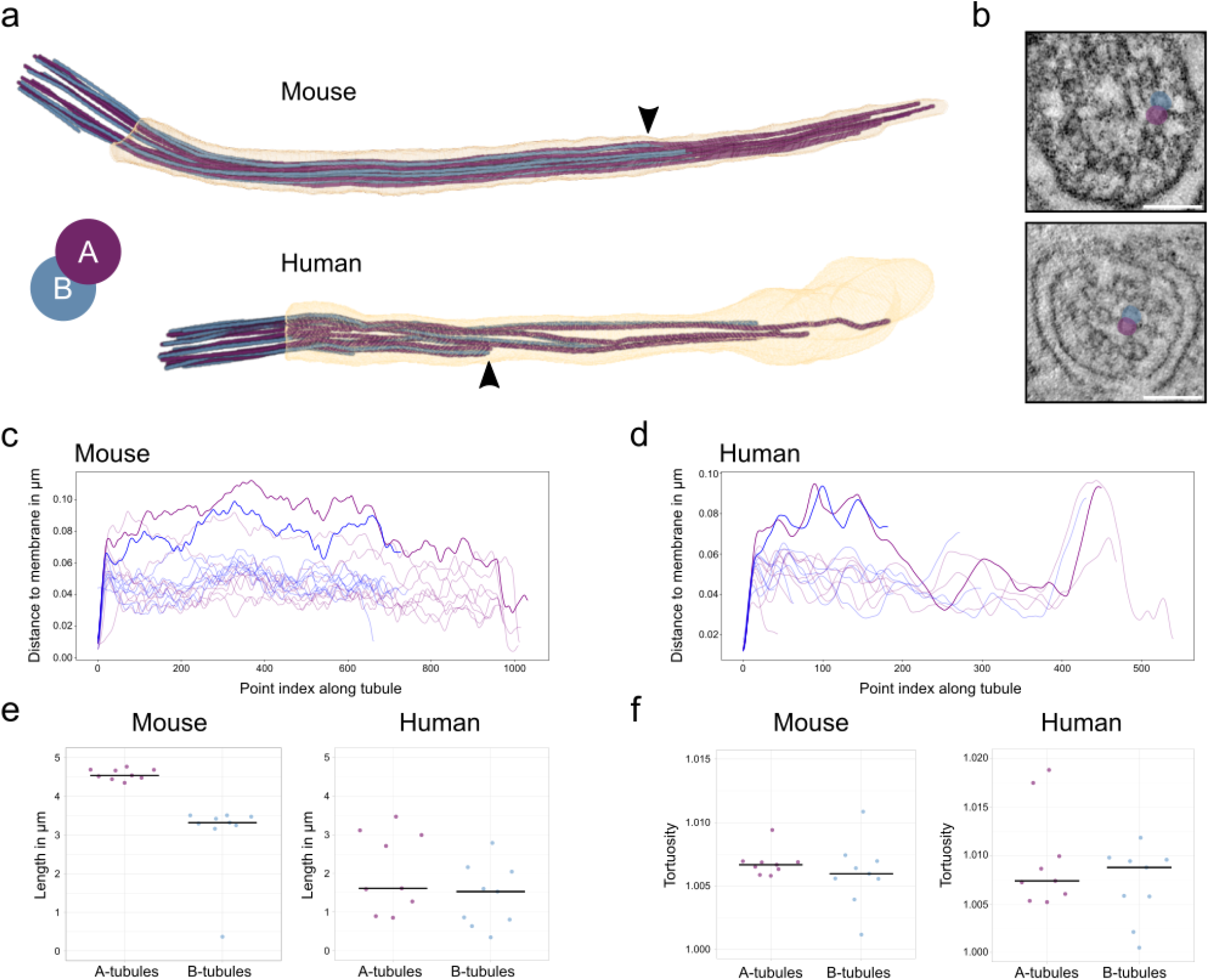
The axoneme structure of primary cilia of mouse and human beta cells. **a)** 3D rendering of mouse and human primary cilia with ciliary membrane (transparent), A-tubules (purple) and B-tubules (blue). **b)** Displacement of microtubule doublets to the center of a human beta cell cilium. A- and B-tubules of the example doublet are highlighted in purple (A) and blue (B). **c)** Distance of the microtubules of the mouse beta cell primary cilium to the ciliary membrane. Doublets with a major transition are highlighted. **d)** Distance of the microtubules of the human beta cell primary cilium to the ciliary membrane. Doublets with a major transition are highlighted. **e)** Length distributions of A- and B-tubules in respective segmentations. **f)** Tortuosity of A- and B-tubules in respective segmentations.

### U-ExM enables quantitative molecular imaging of primary cilia in beta cells and pancreas tissue

Since EM generally does not allow for high throughput imaging and does not readily give molecular information, such as protein identity by antibody labeling and localization, we employed and optimized ultrastructural expansion microscopy (U-ExM) (45) for imaging primary cilia in cultured human beta cells and tissue sections. This method allows for the crosslinking of a biological specimen to a swellable hydropolymer, for its physical expansion and for super-resolution imaging. U-ExM has previously enabled the high-resolution fluorescence imaging of centrioles (45) intraflagellar transport (IFT) trains within motile cilia (46), and of connecting cilia in mouse retina (47). It is therefore ideally suited to collect more quantitative and molecular information on beta cell primary cilia.

We both performed U-ExM on harvested adult pancreas, both by expanding the whole pancreas, and by collecting sections for expansion (Fig. 3a). We further utilized fluorophore conjugated NHS-ester dyes, which bind the amine groups of protein chains and allow for robust fluorescent signals. To this end, we were able to determine cell-types with histology-like precision. Acinar cells were readily identifiable owing to the dark staining of zymogen granules by NHS ester and surprisingly, acetylated tubulin (Fig. 3b). Capillaries were detected by the presence of erythrocytes (Fig. 3b), while the canal-like architecture of ducts was also easily distinguishable (Fig. 3b). Islets were initially identified by large islands without zymogen granules, and confirmed by the presence of insulin staining in the beta cells contained within the islet (Fig. 3c). To investigate potential differences in ciliary frequency, length, or appearance, we stained sectioned pancreas with antibodies against acetylated tubulin and Arl13b, both highly enriched in the cilium (Fig. 3c). Similar to previous reports (48), we found that 80-90% of islets cells were ciliated, including 85% of beta cells as identified by insulin staining (Fig. 3c and d). As in the vEM data primary cilia in the UExM volumes had diverse shapes from straight to very bent (Supp. Fig. 3). We further characterized pancreatic cilia by measuring ciliary length in a semi-automated manner. The cilia of pancreatic ducts were significantly longer (12.41 μm) than the ciliated cells of the islets (approximately 6.8 μm) (Fig. 3e). We detected no appreciable differences in the length of beta cells and other cell types of the islet (Fig. 3e). While imaging islets, we frequently observed the cilia of beta cells making physical contact with other cells, including other beta cells, other cell types in the islets, and surprisingly, acinar cells and capillaries (Fig. 3f, Fig. 4, Supp. Fig. 5). To investigate the ultrastructure of these interactions we used several EM methods. This characterization can be found in the following paragraphs.

**Fig. 3.**
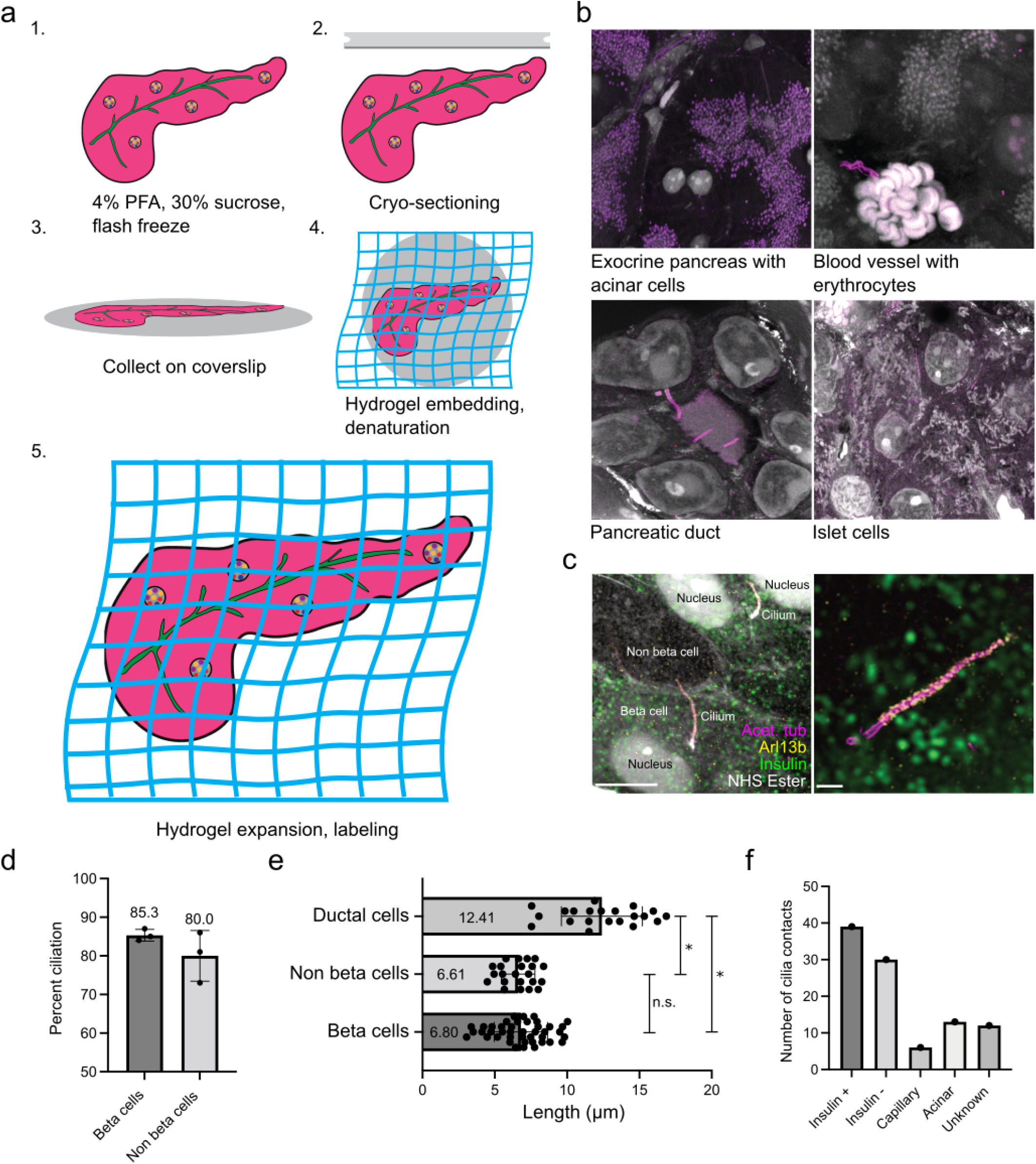
U-ExM of mouse pancreas reveals cilia and cell type identity. **a)** Schematic overview of pancreas U-ExM, including fixation, sectioning, collection, and expansion procedure. **b)** Representative confocal images of expanded pancreatic tissues labeled with antibodies for acetylated tubulin (purple) and with NHS ester for total protein (white). **c)** Higher-magnification images of ciliated islets, containing beta cells, as determined by insulin staining (green), acetylated tubulin (magenta), Arl13b (yellow) and NHS ester. Scale bar, left: 5 μm. Scale bar, right: 1 μm. **d)** Quantification of percent ciliation in beta cells and non beta cells contained within 3 islets from 3 different mice. 50 cells per islet were quantified. **e)** 3D length measurements of the cilium of ductal cilia, non beta cells, and beta cells. Mean length: ductal cilia, 12.41 μm +/- 2.8, non beta cells 6.610 μm +/-1.158, beta cells 6.804 μm +/-1.841. Error reported in standard deviation. Statistical significance measured by unpaired t-test. Ductal cells - beta cells: p <0.0001, ductal cells - non beta cells: p <0.0001, beta cells - non beta cells: p = 0.6546. n = 42 beta cells, 22 non beta cells, 21 ductal cells from 2 pancreas. **f)** Contacts of beta cell cilia to beta- and non-beta islet cells, as well as capillaries, acinar cells and unknown cells.

**Fig. 4.**
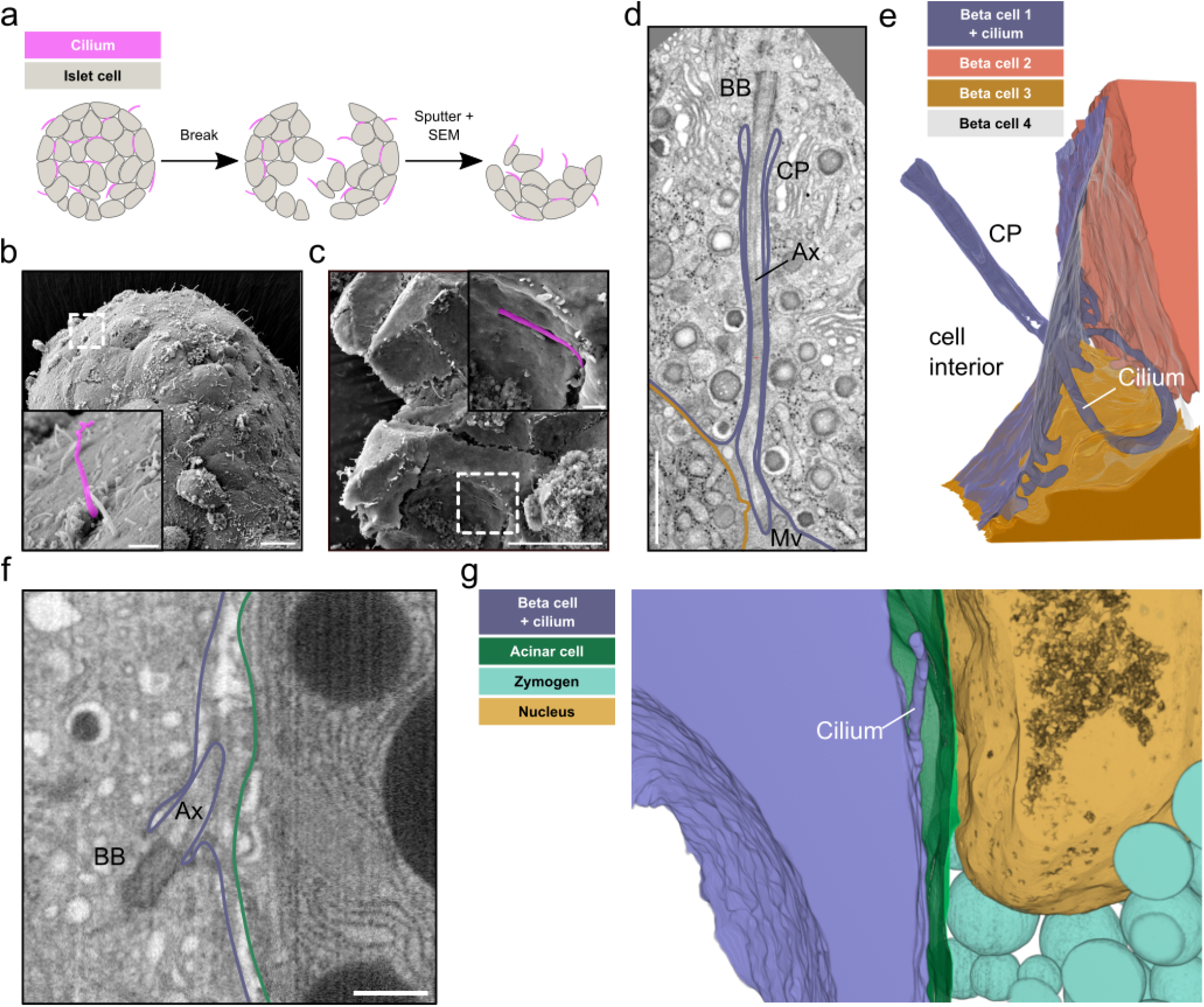
Spatial restriction through ciliary pockets and neighboring cells. **a)** Schematic workflow of breaking dried islets for SEM. **b)** SEM image of the surface of an isolated mouse islet with islet cell primary cilia projecting into the extracellular space. The inset shows a primary cilium highlighted in purple. **c)** SEM image of a broken isolated islet with a primary cilium restricted by the ciliary pocket. The inset shows a magnified view with the primary cilium highlighted in purple. Scale bars overview images: 10 μm, insets: 1 μm. **d)** A single slice of a FIB-SEM volume with a beta cell primary cilium with a long ciliary pocket (CP). The cilium is extending towards the outside where it is surrounded by neighboring cells and microvilli (Mv). Ax: Axoneme, BB: Basal body. Scale bar: 1 μm. **e)** 3D rendering of the segmentation shows the long ciliary pocket and the cilium surrounded by neighboring beta cells. **f)** A single slice of a FIB-SEM volume with a primary cilium of a beta cell on the edge of the islet close to the exocrine tissue. Scale bar: 500 nm. **g)** Segmentation showing the beta cell primary cilium projecting into the extracellular space close to an acinar cell.

### Primary cilia are spatially restricted within long ciliary pockets and by neighboring cells

Complementary to the quantitative data of UExM, EM of beta cell primary cilia deep within isolated islets and pancreas tissue allowed us to assess their ultrastructural features in situ. Previous SEM imaging (30) and our own SEM data of intact isolated islets only allow for imaging their peripheral surfaces. In such cases the cilia appeared as structures extending into an empty extracellular space (Fig. 4b, Supp. Fig. 4a), due to the collagenase-mediated digestion of the surrounding exocrine tissue and extracellular matrix for the isolation of the islets. However, even by SEM it is possible to address a more natural tissue context by “breaking” the islets after critical point drying (Fig. 4a). The islets usually break along the membranes of their cells (Supp. Fig. 4b). This enabled us to visualize islet cell cilia with narrow ciliary pockets and in close contact to neighboring cells (Fig. 4c). To further investigate the spatial restriction of primary cilia we reconstructed cilia and their neighboring cells in our vEM volumes. We could detect a great heterogeneity in the length of the ciliary pockets, which could be several micrometers long (Fig. 4d), but in some cases, such as the mouse beta cell primary cilium depicted in Fig. 1c, extremely short and almost not detectable. The ciliary pockets were in general very narrow, leaving only a few nm between the cilia membrane and the plasma membrane of the respective beta cell (Fig. 4d). Outside of the ciliary pocket, the cilia did not have much more space and were mostly restricted in the extracellular space between adjacent cells (Fig. 4d, Video 3, Video abstract). Depending on their curvature cilia were restricted by one (see Fig. 1a, Video abstract) or multiple islet cells (Fig. 4d, e, Video 3). UExM had revealed a small percentage of beta cell cilia in close proximity to acinar cells and blood vessels. However, our FIB-SEM datasets did not contain these tissues (due to islet isolation) and our ssET of human pancreas data could only provide a limited z-depth and a small field of view. We therefore screened a public FIB-SEM dataset of mouse pancreas (49) including islets, acinar cells and blood vessels for the presence of primary cilia at the edge of the islets close to the vasculature. We could find beta cell cilia at the interface between islet and exocrine tissue (Fig. 4f). 3D segmentation revealed that these cilia, similar to the ones deeper within the islet, were spatially restricted by either exocrine cells (Fig. 4f, g) and/or the extracellular matrix. In some cases we could also observe beta cell primary cilia in close proximity to blood vessels, projecting along their endothelial cells (Supp. Fig. 5). The voxel size of these datasets of 8 nm was not sufficient to fully reconstruct the axonemes. However, the microtubule structure of these cilia resembled the disorganized 9+0 structure described before.

### Primary cilia closely interact with neighboring islet cells and cilia

In our vEM volumes as well as in the UExM images we observed different modes of cilia connectivity: primary cilia contacting neighboring islet cells, and contacts between cilia originating from distinct islet cells. It seems that contacts with neighboring islet cells were mostly the result of the spatial restriction within the islet: once the cilium leaves the pocket and enters the extracellular space it is immediately surrounded by islet cells. Depending on the shape of the cilium we observed interaction with one or more islet cells. For instance, the highly curved cilium shown in Fig. 4d touched three neighboring cells. When investigating these interactions at high resolution we did not detect direct contacts of ciliary membranes with plasma membranes, as there was always a small gap of a few nanometers between them without any electron dense regions characteristic for gap junctions. Furthermore, we observed cilia “pinching” into neighboringcells, bending their plasma membranes (Fig. 5a). This resulted in tunnel-like structures in the neighboring cells, generating almost synapse-like connections. We could observe this behavior also for alpha cell cilia in the FIB-SEM dataset of mouse pancreas (49). In one case an alpha cell cilium was projecting into a neighboring alpha cell ending close to its nucleus (Supp. Fig. 6a and b). In turn, the primary cilium of this second alpha cell was protruding into its adjacent alpha cell (Supp. Fig. 6c and d). Additionally, we detected encounters of cilia of distinct islet cells in the extracellular space (Fig. 5b). In most cases cilia projected into the same region surrounded by microvilli and their tips got into close proximity to each other. We also observed cilia from a double-ciliated beta cell meeting other cilia on opposite sides of the cell (Supp. Fig. 7). Moreover, in one instance we even found cilia from distinct islet cells meeting in the same ciliary pocket of a beta cell (Fig. 5c), although in this case we could not clearly determine the cell origin of one of these cilia. This ciliary pocket was relatively wide compared to the other pockets we observed and it also contained large vesicular structures.

**Fig. 5.**
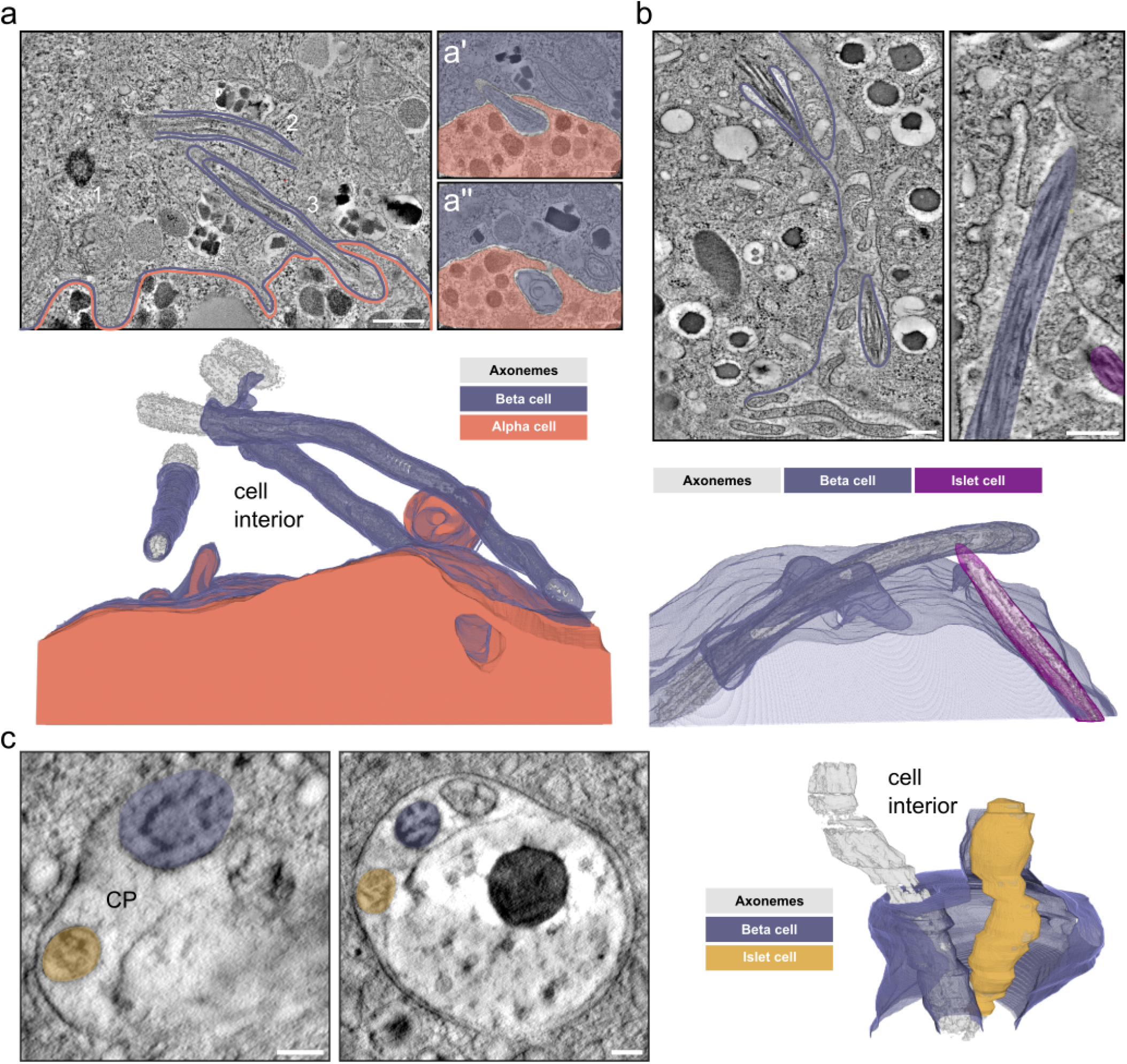
Cilia-cell and cilia-cilia interaction. **a)** Single tomographic slices of primary cilia in human islets. There are 3 cilia originating in a beta cell (purple). The neighboring alpha cell is outlined in orange. In magnified views of distant slices through the tomogram (a’ and a”) the end of cilium 2 is visible protruding and pushing into the neighboring alpha cell (orange). Axonemes, basal bodies and centrioles are rendered in gray. Scale bars: 500 nm and 100 nm. **b)** Single tomographic slices through mouse beta cells with 2 primary cilia (purple and magenta) originating from a beta cell (purple) and another islet cell (magenta) meeting in the space between cells. The side view shows the axonemes of the cilia in close proximity. The 3D view shows the segmentation of ciliary and plasma membranes together with the axonemes. Scale bars: 200 nm. **c)** Tomographic slices through an ssET volume showing a large ciliary pocket that is shared by a primary cilium originating from the beta cell (purple) and a primary cilium of unknown origin (yellow). The 3D rendering shows the interaction of the two cilia. Axonemes in gray. Scale bars: 100 nm.

### Beta cells can contain multiple cilia

Previous studies reported the presence of two primary cilia in a few beta cells (26; 30). We could make similar observations in mouse pancreas tissue by UExM and found beta cells with two primary cilia (Fig. 6a). In all UExM images of mouse beta cell primary cilia we could observe a basal body and a daughter centriole for each cilium in double-ciliated cells, indicating that cilia were not originating from daughter centrioles in mouse beta cells. Also, the basal bodies had variable distances to each other but the axonemes of both cilia were in close proximity to each other. By vEM, we observed human beta cells with three primary cilia Fig. 5a. The cilia appeared to have similar structural features as those from single-ciliated beta cells with the same disorganized axonemes. Furthermore, in FIB-SEM data of mouse islets we observed beta cells with two primary cilia (Fig. 6b, c). Here, we could discriminate between cilia sharing one ciliary pocket (Fig. 6b) and multiple cilia in separate pockets (Fig. 6c). In the latter case the cilia had very long ciliary pockets and left the cell on opposite sides, meeting other cilia as described in the previous paragraph (Supp. Fig. 7). Notably, we did not detect beta cells with three primary cilia in our mouse datasets.

**Fig. 6.**
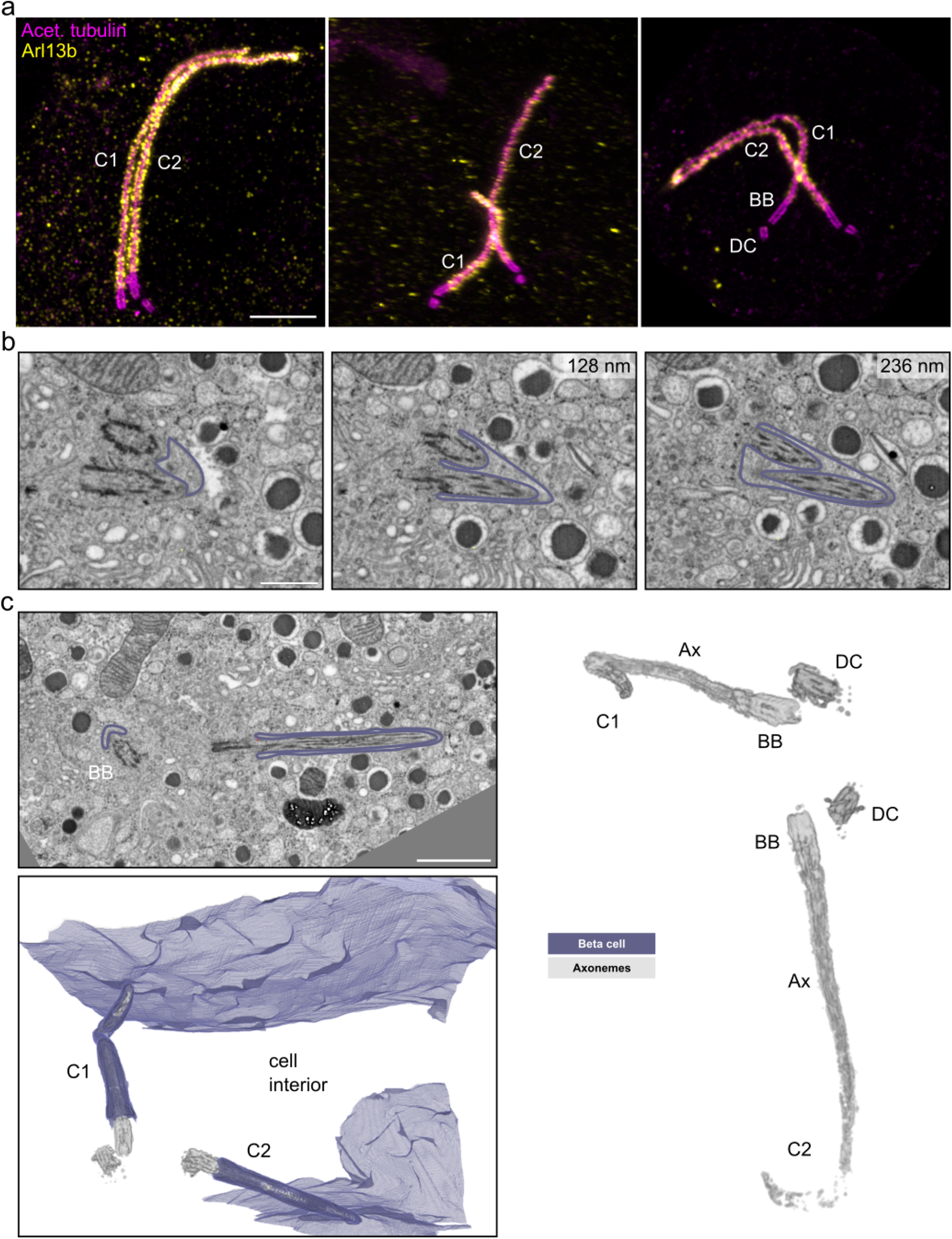
Multiple cilia in beta cells. **a)** Expanded multiple islet cell cilia stained by acetylated tubulin (magenta) and Arl13b (yellow). Scale bar: 1 μm. **b)** Single FIB-SEM slices showing a beta cell containing two primary cilia sharing one ciliary pocket. The membrane is outlined in purple. Z-distances from the first image are indicated in the upper right of the following slices. Scale bar: 500 nm. **c)** FIB-SEM data with a beta cell containing two primary cilia (C1 and C2) with distinct ciliary pockets. The cilia are pointing in opposite directions. The raw FIB-SEM slice shows the two cilia with membranes outlined in purple. Scale bar: 1 μm. The 3D rendering shows the plasma membrane in purple and the cilia microtubule structures in gray. The rendering on the right shows basal bodies (BB), daughter centrioles (DC) and axonemes (Ax) in gray.

### Islet cell primary cilia connect to the pancreatic innervation

Research has so far focused on the role of islet cell cilia in paracrine signaling between endocrine cells. Recently, however, primary cilia have been shown to form synapses with neurons in brain tissue (50). While screening the aforementioned 8 nm isotropic FIB-SEM dataset of mouse pancreas tissue (49) we found several nerve fibers contacting the islets (Fig. 7a). These fibers showed classical structural features of neuronal axons, being very long, containing cables of parallel microtubules and synaptic vesicles (Fig. 7a). They usually reached the edge of the islet and occasionally split into few fibers terminating deeper in the islet. They contacted the plasma membranes of islet cells including alpha, beta and delta cells. Surprisingly, we detected primary cilia (as identified by the presence of basal body, daughter centriole and axoneme) of beta and alpha cells projecting towards these nerve fibers and contacting them (Fig. 7b,c, Supp. Fig. 8, Video 4). On some of these contacts we could observe structural features of presynapses on the side of the neurons: synaptic vesicles close to these contacts and possibly synaptic vesicles close to these contacts and possibly appearance of synaptic vesicle exocytosis (Fig. 7b’, b”, Video 4). We observed these contacts being formed by both beta cell as well as alpha cell cilia, as the axons traversed through the islet (Fig. 7c). In this case the primary cilia of a beta cell and the adjacent alpha cell connected with the same nerve fiber. In most cases cilia were contacting axons with their lateral side. However, we could also observe a case of an alpha cilium touching an axon with its tip (Supp. Fig. 8).

**Fig. 7.**
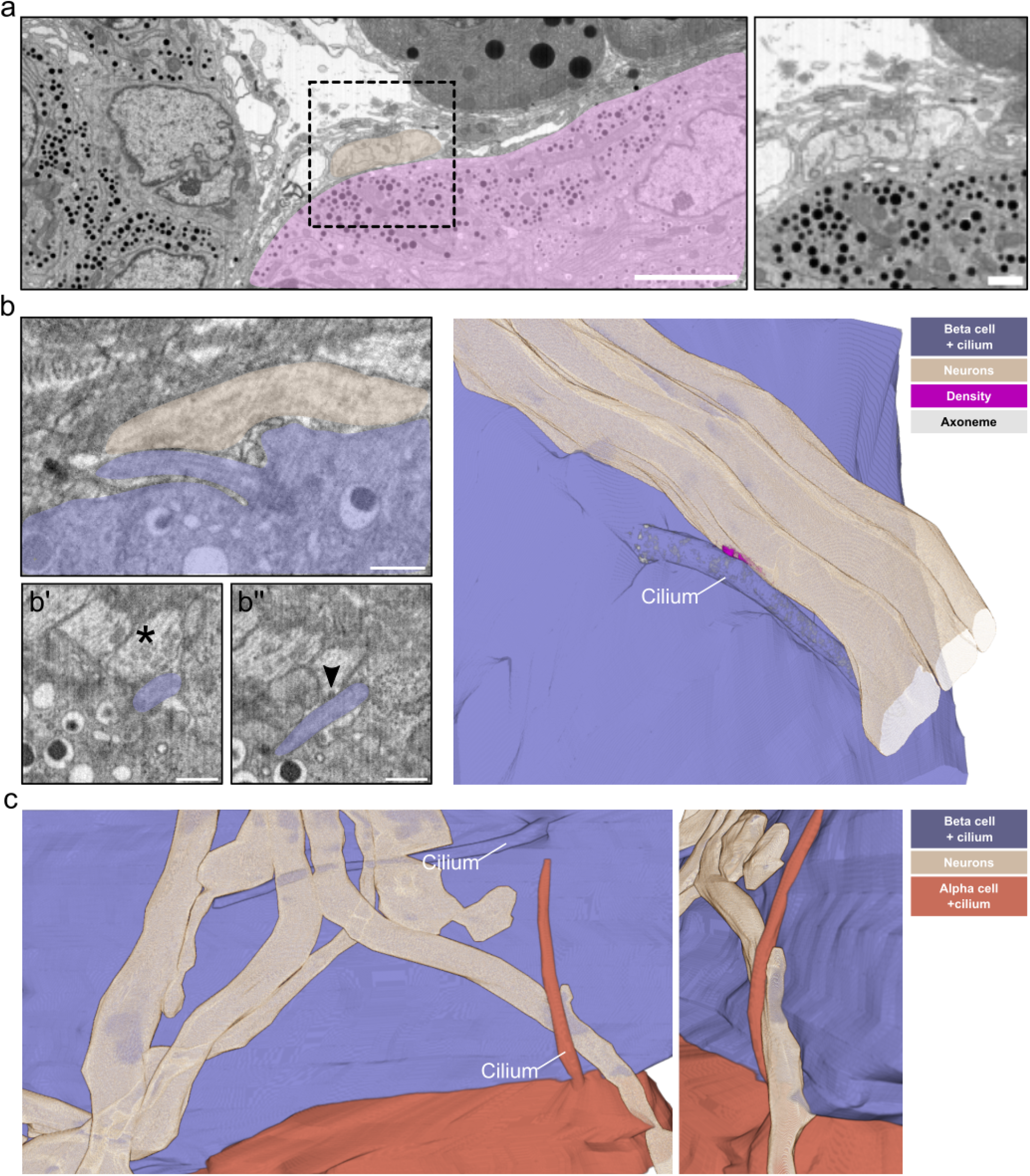
Islet cell primary cilia - neuron connections. **a)** Slices through a FIB-SEM volume showing axons (beige) connecting to a pancreatic islet (magenta). Scale bar: 5 μm. Next is the magnification of the boxed area showing the different neurons in contact with an alpha cell. Scale bar: 1 μm. **b)** A single slice of a FIB-SEM volume with an axon (beige) and beta cell with a primary cilium (purple) in contact with each other. Scale bar: 1 μm. The asterisk in b’ highlights synaptic vesicles close to the cilium-neuron contact. The arrowhead in b” points to a likely fusion event of a synaptic vesicle at the synaptic density. Scale bars: 500 nm. The 3D rendering shows the contact of a primary cilium with one of the axons, including a segmentation of the synaptic density. **c)** 3D rendering of primary cilia from a beta and an alpha cell in contact with the same axon. The smaller image shows a tilted magnified view of the alpha cell cilium-axon interaction.

## Discussion

We provide here the first in-depth volumetric characterization of primary cilia within mouse and human tissue by applying vEM and U-ExM, thus resolving the structural organization of the axoneme and giving a comprehensive overview on cilia interaction and connectivity. High resolution 3D data have so far only been available for primary cilia of monolayer cells facing the cell culture medium (23; 22). Our data allow us to resolve the structure of the axoneme as well as the spatial interaction of islet beta cell primary cilia in their native environment. We can show that the axoneme is not fully maintained throughout the whole length of the cilium in mouse and human beta cell cilia. Together with neurons, glia, and cultured kidney cells, this marks another cell type in which a continuous 9+0 organization is the exception, not the rule, and strongly suggests that most primary cilia deviate from this organization. In the reconstructed human beta cell cilium the length of microtubules of the axoneme varied much more compared to the mouse beta cell cilium. This might be explained by the old age of the human beta cells. However, these values will need to be supported by data of more cells in the future. Notably, the primary cilium has been shown to be one of the oldest organelles in beta cells (51), which might explain a stronger dis-organization as the cells age. We further show the absence of a central microtubule pair along the full length of the cilium. Instead, some of the outer microtubule doublets/singlets are displaced towards the center of the cilium, which explains previous observations of central microtubule doublets in 2D sections (35). While these data do not fully rule out the possibility of cilia movements, they indicate that the previously observed phenomena (35) cannot be mediated by the microtubule organization of the axoneme. Notably, primary cilia have been demonstrated to move by Brownian motion and processes mediated by the actin cytoskeleton surrounding the basal body (52) - factors that might at least contribute to beta cell cilia motility.

Imaging beta cell primary cilia in their in situ environment -isolated pancreatic islets and pancreas - enabled us to investigate their interactions within the tissue context. We found that beta cell cilia are spatially restricted within the islets of Langerhans by their own ciliary pockets and the neighboring cells, leaving only a few nanometers of freedom. This spatial restriction resulted in the formation of connections of cilia with neighboring cells. Depending on the shape of the cilia they could reach several islet cells. If these cilia exert special functions within the islet remains to be investigated. Furthermore, cilia formed synaptic-like structures pinching neighboring cells. Cilia can convey the signal of developmental factors, such as WNT (53) and hedgehog (54). Recently, it has been shown that WNT4 regulates the physiology of mouse adult beta cells (55). Yet, the molecular factors involved in the establishment of these connections and the timing of their formation are still unknown. The signals transmitted or received via the cilium are also largely elusive. Beta cell cilia have been shown to be involved in GABA signaling (56). Furthermore, they express the insulin (33), somatostatin (57; 58), and various G protein-coupled receptors (48). If some of these pathways are especially prominent at cilia-islet cell connections remains to be investigated. Even more surprising to us was the observation of interacting cilia from distinct islet cells. They met not only in the extracellular space, but even in shared ciliary pockets. These phenomena could be the result of the special islet architecture (5; 6). However, destruction of beta cell cilia by knocking out of *IFT88* (32) resulted in loss of beta cell polarity. Thus, we believe that our observations are most likely not random events, but reflect specific signaling events that attract cilia to each other. The molecular pathways and factors supporting these cilium-cilium interactions are also unknown. However, cilia have been described to support cell connectivity in the brain (59; 60) and might play a similar role within pancreatic islets. The evidence that beta cells can establish connections with neighboring cells through multiple cilia further support the idea of cilia being relevant for paracrine communication. It is known that beta cells are heterogeneous and that so-called “hub” or “leader” beta cells play a role in islet calcium signaling (61; 62). Further work shall focus on the functional characterization of these multiciliated beta cells and on their role within the islet.

In each example of multiple ciliated beta cells, two complete copies of the centrosome were present, with the cilium emanating from the mother centriole, and two daughter centrioles present in the vicinity. This feature is in contrast to the multiple cilia occasionally observed in cultured cells, where a cilium emerges from the daughter centriole or bifurcates near the base. Cilia emerging from two distinct mother centrioles reflects a controlled and distinct process of centrosome duplication to allow the formation of two complete independent signaling entities.

Recent publications investigating the role of primary cilia in adult human brain cortex (63) and mouse visual cortex (64) have revealed ultrastructural differences in the cilium pertaining to cell type, specifically in the ciliary pocket. Interestingly, the embedding of beta cell primary cilia deep into ciliary pockets closely resembles features observed in astrocytes (64; 63). These cells have a well described role in maintaining connectivity with endothelial cells at the blood brain barrier (65). We speculate that physical contacts between beta cell cilia and endothelial cells modulate their reciprocal signaling. Notably, we found beta cell cilia with very long and very short ciliary pockets. In neuronal tissue cilia with long pockets do not reach as many cells as cilia with short pockets (64; 63). However, cilia with long pockets have more surface area for potential autocrine signaling. Furthermore, the ciliary pocket is a hub for membrane trafficking such as endocytosis (66), which could be enhanced in cilia with long pockets.

Finally, the observation of synapses between islet cell cilia and innervating axons points to signaling functions of primary cilia beyond the local islet context. To our knowledge these kinds of connections (serotonergic cilia-axon synapses) have only been described in the brain (50). Thus, our reconstructions are the first to document this phenomenon elsewhere in the body. As among neurons, synaptic-like contacts among beta or alpha cells through their cilia may enable a more direct access to the inner parts of the receiving cell, especially to its nucleus. In the brain serotonergic axon-cilium synapses cause chromatin alterations with epigenetic changes. We did not yet identify neurotransmitters involved in synaptic-like contacts of beta cell cilia but future studies in this direction will likely provide insights into the physiology of islet cells in health and diabetes. In particular, we speculate that such ciliary synaptic-like contacts represent a privileged site for the response of islet cells to sympathetic/parasympathetic (67) and paracrine signals. Furthermore, recent work postulates that the close vicinity of cilia to synapses enables them to detect synaptic neurotransmitters (64; 63). This could be the case also for islet cell cilia touching axons without the formation of cilia-axon synapses. In summary, we provide novel structural insights into the role of primary cilia as a key factor in islet cell connectivity and signaling. Our work opens new research questions on the pathways for establishing cilia connectivity and the molecular factors involved in ciliary signaling.

## Supporting information

Video abstract

## Data and software availability

The raw FIB-SEM dataset we used for segmentation of the mouse beta cell cilium axoneme can be found at: https://openorganelle.janelia.org/datasets/jrc_mus-pancreas-3

The raw FIB-SEM datasets used for investigation of cilia restriction and interaction can be found at: https://openorganelle.janelia.org/datasets/jrc_mus-pancreas-1 and https://openorganelle.janelia.org/datasets/jrc_mus-pancreas-2

## ACKNOWLEDGEMENTS

We thank the electron microscopy facility of MPI-CBG for their services, as well as the National Facility for Light Imaging at Human Technopole. We thank Katja Pfriem for administrative assistance. We thank members of the PLID, especially Lisa-Marie Kretschmar, for valuable feedback. We thank Olof Idevall-Hagren (Uppsala University, Sweden) for critical comments on the manuscript. We thank Isabel Espinosa-Medina, Wei-Ping Li, Zhiyuan Lu, Wei Qiu, Eric Trautman, Stephan Preibisch, Davis Bennett, Yurii Zubov, Rebecca Vorimo, Aubrey Weigel, and the CellMap Project Team (all HHMI/Janelia) for generating and sharing the FIB-SEM dataset of P7 mouse pancreas. This work was supported with funds to M Solimena from the German Center for Diabetes Research (DZD e.V.) by the German Ministry for Education and Research (BMBF), from the German-Israeli Foundation for Scientific Research and Development (GIF) (grant I-1429-201.2/2017), from the German Research Foundation (DFG) jointly with the Agence nationale de la recherche (ANR) (grant SO 818/6-1), and from the Innovative Medicines Initiative 2 Joint Undertaking under grant agreements no. 115881 (RHAPSODY) and no. 115797 (INNODIA), which include financial contributions from the European Union’s Framework Programme Horizon 2020, EFPIA and the Swiss State Secretariat for Education, Research and Innovation (SERI) under contract number 16.0097, as well as JDRF International and The Leona M. and Harry B. Helmsley Charitable Trust. G Pigino is supported by Human Technopole and the European Research Council (ERC) under the European Union’s Horizon 2020 research and innovation program (grant agreement no. 819826). AM was the recipient of a MeDDrive grant from the Carl Gustav Carus Faculty of Medicine at TU Dresden. NK was the recipient of EMBO Fellowship ALTF 537-2021. CSX, SP and HFH are supported by the Howard Hughes Medical Institute. M Seliskar was supported by the International Federation of Medical Students Association (IFMSA). DS was supported by the Research Experience Program of TU Dresden. DS was funded by HELMHOLTZ IMAGING, a platform of the Helmholtz Information & Data Science Incubator.

C.S.X. and H.F.H. are the inventors of a US patent assigned to HHMI for the enhanced FIB-SEM systems used in this work: Xu, C.S., Hayworth, K.J., and Hess, H.F. (2020) Enhanced FIB-SEM systems for large-volume 3D imaging. US Patent 10,600,615, 24 Mar. 2020. The other authors declare no competing interests. Author contributions: Project design and supervision: A.M., N.K., G.P., M.Solimena; Data acquisition and analysis: A.M., N.K., S.P., L.E.G.G., D.S., O.T.; M. Seliskar, D.S., C.S.X.; Patient recruitment and surgery: J.W. and M.D.; Sample collection and processing: E.S., N.K., D.F., S.K., C.M., H.M.; Writing-Original Draft: A.M., N.K., G.P., M.Solimena; Writing-Review and Editing: All coauthors

## Materials and Methods

### Islet isolation and culture

Isolation of islets of Langerhans of 9-week-old C57BL/6 mice was performed as previously described (68). Islets were cultured overnight in standard culture media (Roswell Park Memorial Institute 1640 [Gibco] with 10% FBS, 20 mM Hepes, and 100 U/ml each of penicillin and streptomycin) with 5.5 mM glucose. Before processing by HPF, the islets were incubated for 1 h in Krebs–Ringer buffer containing either 3.3 mM or 16.7 mM glucose. All animal experiments were conducted according to guidelines of the Federation of European Laboratory Animal Science Associations (FELASA) and are covered by respective licenses for those experiments from the local authorities. Facilities for animal keeping and husbandry are certified and available with direct access at the Dresden campus (including facilities at Paul Langerhans Institute Dresden and Max Planck Institute for Molecular Cell Biology and Genetics). Licenses for animal experiments are approved by the State Directorate Saxony under license number DD24.1-5131/450/6.

### Preparation of pancreatic specimens of living donors for vEM

Pancreas specimens of a cohort of patients from the University Hospital Carl Gustav Carus described in (41) were examined by a certified pathologist after resection. The samples were cut into cubes with a side-length of max. 1 mm and fixed with 4% glutaraldehyde in sodium phosphate buffer immediately after dissection. Samples were block-contrasted with 1% osmiumtetroxide followed by 1% uranylacetate. After dehydration samples were embedded in epoxy resin (Embed 812, Science Services). Ultrathin sections were cut with a Leica U6 ultramicrotome (Leica Microsystems) and post-stained with uranylacetate and leadcitrate. Grids were screened on a JEM 1400 Plus with a Jeol Ruby camera. The sample with the best preservation of ultrastructure (from a patient with impaired glucose tolerance) was used for cutting serial sections for ssET.

### High pressure freezing, freeze substitution and embedding

Isolated islets were kept in culture overnight and frozen with a Leica EMpact 2 or EM ICE high-pressure freezer (Leica Microsystems). They were kept in liquid nitrogen until freeze substitution. Freeze substitution was performed according to a previously published standard contrast protocol (69) or according to the following high contrast protocol (70; 28): first the samples were substituted in 2% osmiumtetroxide, 1% uranylacetate, 0.5% glutaraldehyde, 5% H_2_O (according to (71)) in acetone with 1% methanol at −90°C for 24 hours. The temperature was raised to 0°C over 15 hours followed by 4 washes with 100% acetone for 15 min each and an increase in temperature to +22°C. Afterwards the samples were incubated in 0.2% thiocarbohydrazide in 80% methanol at RT for 60 min followed by 6×10 min washes with 100% acetone. The specimens were stained with 2% osmiumtetroxide in acetone at RT for 60 min followed by incubation in 1% uranylacetate in acetone + 10% methanol in the dark at RT for 60 min. After 4 washes in acetone for 15 min each they were infiltrated with increasing concentrations of Durcupan resin in acetone followed by incubation in pure Durcupan and polymerization at 60°C. For quality control the blocks were sectioned with a Leica LC6 ultramicrotome (Leica microsystems) and 300 nm sections were put on slot grids containing a Formvar film. Tilt series ranging from −63° to +63° were acquired with a F30 electron microscope (Thermo Fisher) and reconstructed with the IMOD software package (72).

### Serial sectioning and electron tomography

Blocks were sectioned with a Leica LC6 ultramicrotome (Leica Microsystems), and 300 nm serial sections were put on slot grids containing a Formvar film. Sections were contrasted with 2% uranylacetate in methanol followed by 1% leadcitrate in H_2_O. Tilt series ranging from −63° to +63° were acquired with a F30 EM (Thermo Fisher Scientific). The tomograms were reconstructed and joined with the IMOD software package reconstructed with the IMOD software package (72).

### FIB-SEM imaging

Multiple Durcupan embedded isolated islets samples were each first mounted to the top of a 1 mm copper post which was in contact with the metal-stained sample for better charge dissipation. The region of interest (ROI) was identified by x-ray tomography data obtained with a Zeiss Versa XRM-510 and optical inspection under a microtome. These vertical sample posts were each trimmed to its defined ROI using a Leica UC7 ultramicrotome. This sample preparation methodology was developed for precise ROI targeting and improved image acquisition, as previously described (73). Before FIB milling, a thin layer of conductive material of 10 nm gold followed by 100 nm carbon was coated on each sample with a Gatan 682 Precision Etching and Coating System with the following coating parameters: 6 keV, 200 nA on both argon gas plasma sources, 10 rpm sample rotation with 45° tilt. After coating, each sample was imaged with a customized, enhanced FIB-SEM consisting of a Zeiss Capella FIB column mounted at 90° onto a Zeiss Merlin SEM as previously described (74; 29). Each block face was imaged by a 140 pA electron beam with 0.9 keV landing energy at 200 kHz. The x-y pixel size was set at 4 nm. A subsequently applied focused Ga+ beam of 15 nA at 30 keV strafed across the top surface and ablated away 4 nm of the surface. The newly exposed surface was then imaged again. The ablation–imaging cycle continued about once every 3–4 min for several days up to two weeks to complete the FIB-SEM imaging of one sample. The sequence of acquired images formed a raw imaged volume, followed by post-processing of image registration and alignment using a Scale Invariant Feature Transform–based algorithm. The actual z-step was estimated by the changes of SEM working distance and FIB milling position. The image stacks were rescaled to form 4 × 4 × 4 nm isotropic voxels, which can be viewed in any arbitrary orientations. In some cases FIB-SEM stacks were denoised with the DenoisEM plugin (75) in FIJI (76).

### Segmentation and Analysis

Membranes were manually segmented in Microscopy Image Browser (MIB) (77). Rough segmentations of the axonemes were performed also in MIB by local thresholding. Segmentation of microtubules was performed by skeleton tracing in Knossos (78). All segmentation data were centrally loaded into the CellSketch viewer within the Album software (79) according to our recently published protocol (80). This workflow was also used to generate data on microtubule lengths which were plotted with PlotsOf-Data (81). Segmentation masks were rendered with Blender (www.blender.org) according to our protocol or ORS Dragonfly (www.theobjects.com/dragonfly/index.html). For the rendering in Fig. 1c and Supp. Video x we segmented the cells organelles with Autocontext in ilastik (82).

### SEM imaging

Isolated mouse islets were fixed in 4% formaldehyde in 100mM phosphate buffer, followed by post-fixation in modified Karnovsky (2% glutaraldehyde and 2% formaldehyde in 100mM phosphate buffer). The samples were washed 2 × 5 min with PBS and 3 × 5 min with bi-distilled water and post-fixed in 1% osmium tetroxide in water for 2 h on ice, followed by washes in water (6x 5 min), dehydration in a graded series of ethanol/water mixtures up to pure ethanol (30%, 50%, 7%, 96% 15 min each, and 3 × 100% on molecular sieve, 30 min each) and critical point drying using a Leica CPD300. Dried samples (complete islets or broken specimens) were mounted on 12 mm aluminum stubs using conductive carbon tabs. To increase contrast and conductivity, samples were sputter coated with gold (BAL-TEC SCD 050 sputter coater, settings: 60 sec, with 60 mA, at 5 cm working distance). Finally, samples were imaged with a JSM 7500F scanning electron microscope (JEOL, Freising, Germany) running at 5kV (using the lower SE-detector and working distances between 3 and 8mm).

### U-ExM of mouse pancreas

Whole mouse pancreas were harvested from adult C57BL/6 mice and fixed overnight in 4% PFA at 4°C. U-ExM was performed on mouse pancreas by two different methods described below, those being whole tissue expansion and expansion of sectioned tissue. 2% acrylamide, 1.4

### Whole tissue expansion, adapted from (83)

After overnight 4% PFA fixation, whole pancreas were incubated in 2% acrylamide, 1.4% formaldehyde for 72 hrs in 1.5 mL tubes. After the crosslinking prevention, the acrylamide / formaldehyde solution was removed, and 500 μL of inactivated monomer solution (containing .1% triton) was added to submerge the pancreas, and incubated overnight at 4°C. The following day, the inactivated monomer solution was removed. 500 μL of monomer solution supplemented with APS and TEMED (.05% final concentration), as well as .1% triton. The samples were incubated for 1 hr at 4°C, followed by 2 hrs at 37°C in a 1.5 mL tube. Following gelation, the samples were removed from the 1.5 mL tube with a fishing hook, and placed in a 6-well plate containing 1 mL of denaturation buffer. Samples were shaken at room temperature for 10 minutes, the 6 well plate was wrapped with parafilm, and incubated for 72 hrs at 70°C. Following denaturation, the gels were added to a 145 mm dish containing ddHH20 for 30 minutes to initiate expansion. The gels were washed 1X for 30 minutes with ddH20, and incubated overnight in ddH20 to reach full expansion. The following day, the gels were shrunk by washing the sample 2X in 1X PBS for 15 minutes. Upon shrinking, the gels were incubated with the desired primary antibodies in PBS-BSA 2%, for 72 hrs at 37°C while shaking. After primary antibody labeling, the gels were washed 3X for 15 minutes, and were then incubated with secondary antibodies for 48 hrs at 37°C while shaking. The primary antibodies used in this study were mouse-anti-insulin (Sigma), mouse-anti-acetylated tubulin (Sigma), rabbit-anti-Arl13b (Proteintech). For NHS Ester stained gels, a stock concentration of NHS Ester 405 of 1 mg / mL was diluted to a 20 μg / mL working solution in 1X PBS. Samples were incubated for 3 hrs at room temperature while gently shaking. Samples were then washed in 1X PBS, 5 times by 5 minutes. After secondary antibody washing and / or NHS staining, gels were washed 2X for 30 minutes with ddH20, and then incubated in water until imaging.

### Expansion of sectioned tissue, adapted from (47)

After overnight 4% PFA fixation, samples were washed 4X, for 5 min, in 1X PBS. Whole pancreas were then incubated in 30% sucrose solution for 12 hrs at 4°C. Immediately afterwards, the pancreas were incubated in 15% sucrose for 12 hrs at 4°C. The following morning, the pancreas were removed from 15% sucrose and excess solution was blotted away with filter paper. A cryomold was subsequently filled with a thin layer of OCT, the pancreas were placed inside of a cryomold. The cryomold was placed on dry ice, and was filled with OCT. Upon freezing of the OCT, the cryomold was collected and immediately transferred to a Leica CM1950 cryostat for tissue sectioning. Mouse pancreas were transversely sectioned in 20 μm increments, and individually collected on poly-d-lysine (PDL) coated, 12 mm coverslips. Sections were stored in a sealed box at −80°C overnight, and expansion proceeded the next day as described in section 8.

### Fluorescence microscopy of U-ExM gels

Expanded gel quarters were placed in a dish and cut into an approximately 1.5 cm × 1.5 cm square piece. The gel piece was then placed on a 24 mm coverslip, and added to a 35 mm imaging chamber. The chamber was placed on a Zeiss LSM980-NLO Airyscan 2 confocal microscope, and the side containing cells or tissue was identified in widefield mode. Upon correct siding, the gel was removed from the chamber, and gently blotted to remove excess water, and placed cell or tissue side facing down on PDL coated coverslip and added back to the imaging chamber. Z stacks were acquired in Airyscan mode with maximum resolution selected, a 1.4NA, 63X, oil immersion objective, and a z-step size of 0.15 μm.

### U-ExM image quantification

Z-stacks were loaded in Fiji. For length measurements, the simple neurite tracer plugin was utilized to trace the tubulin fluorescence signal of the cilium from the bottom of the basal body in a semi-automated manner, as previously described for IFT trains (46).

## Supplementary Note 1: Supplementary Figures

**Supplementary Figure 1.**
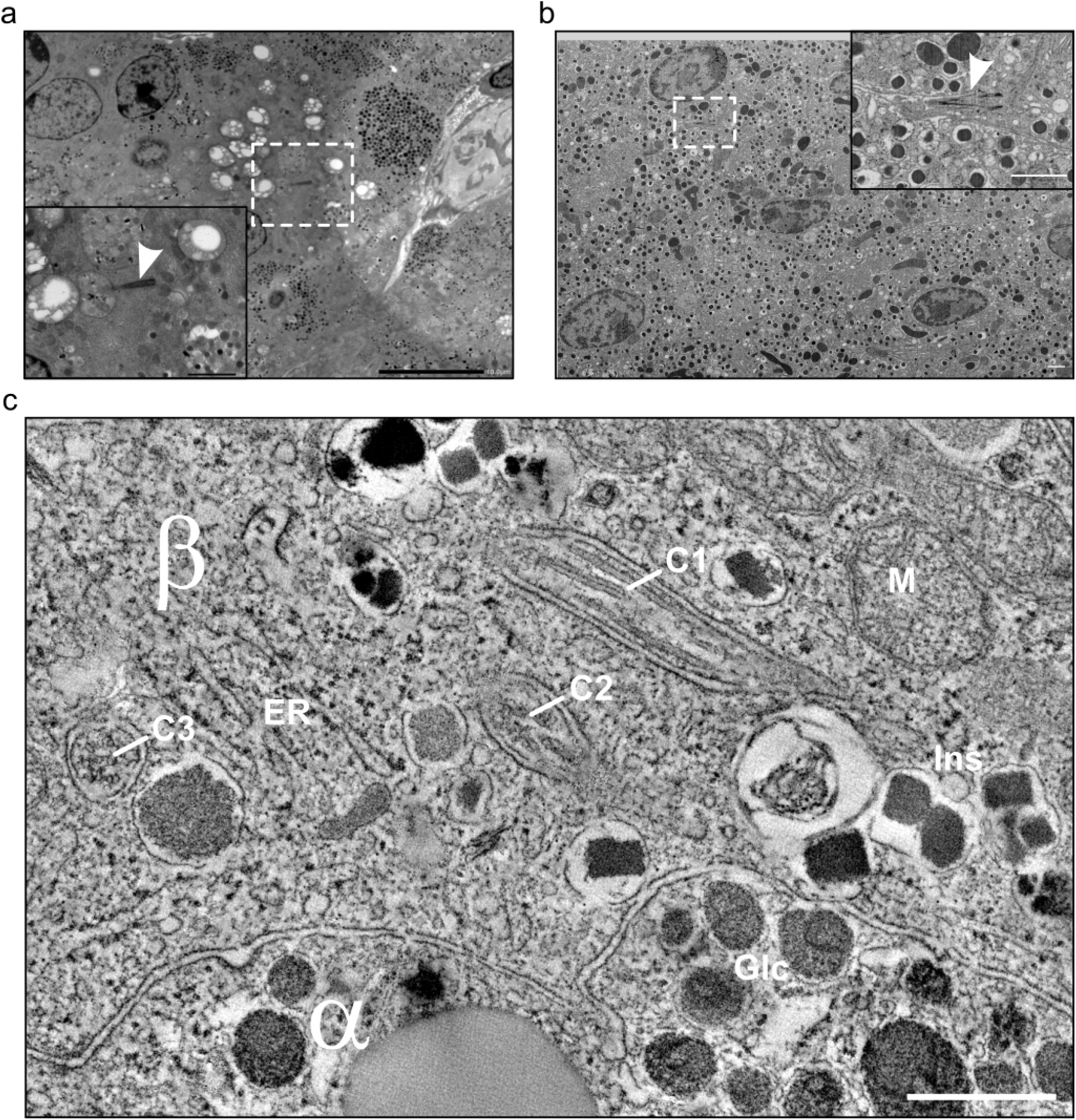
Identification of primary cilia in ssET and FIB-SEM. **a)** TEM overview of a 300 nm thick section of human pancreas. Inset shows a primary cilium identified by the axoneme structure. Scale bars: 10 and 1 μm. **b)** FIB-SEM slice of mouse pancreatic islets. Inset shows a primary cilium. Scale bars: 1 μm. **c)** Single slice of a tomogram with a beta cell (β) with insulin secretory granules (Ins), mitochondria (M), endoplasmic reticulum (ER) and cilia (C1-3). The neighboring alpha cell (α) with glucagon granules (Glc). Scale bar: 1 μm.

**Supplementary Figure 2.**
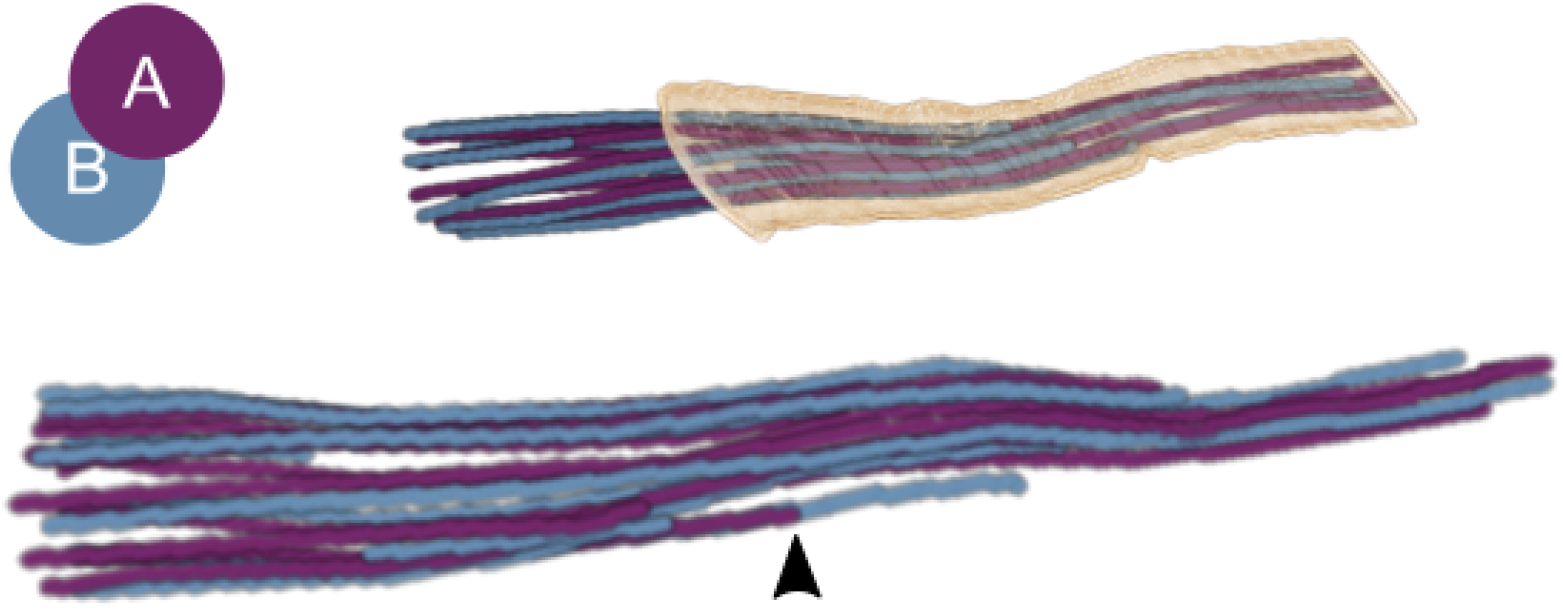
Reconstruction of incomplete human beta cell primary cilium with longer B-tubules. 3D rendering of A-(purple) and B-tubules (blue) with ciliary membrane (light orange). Below is a magnified rotated rendering with an arrowhead pointing to the earlier determination of the A-tubule of the specific microtubule doublet.

**Supplementary Figure 3.**
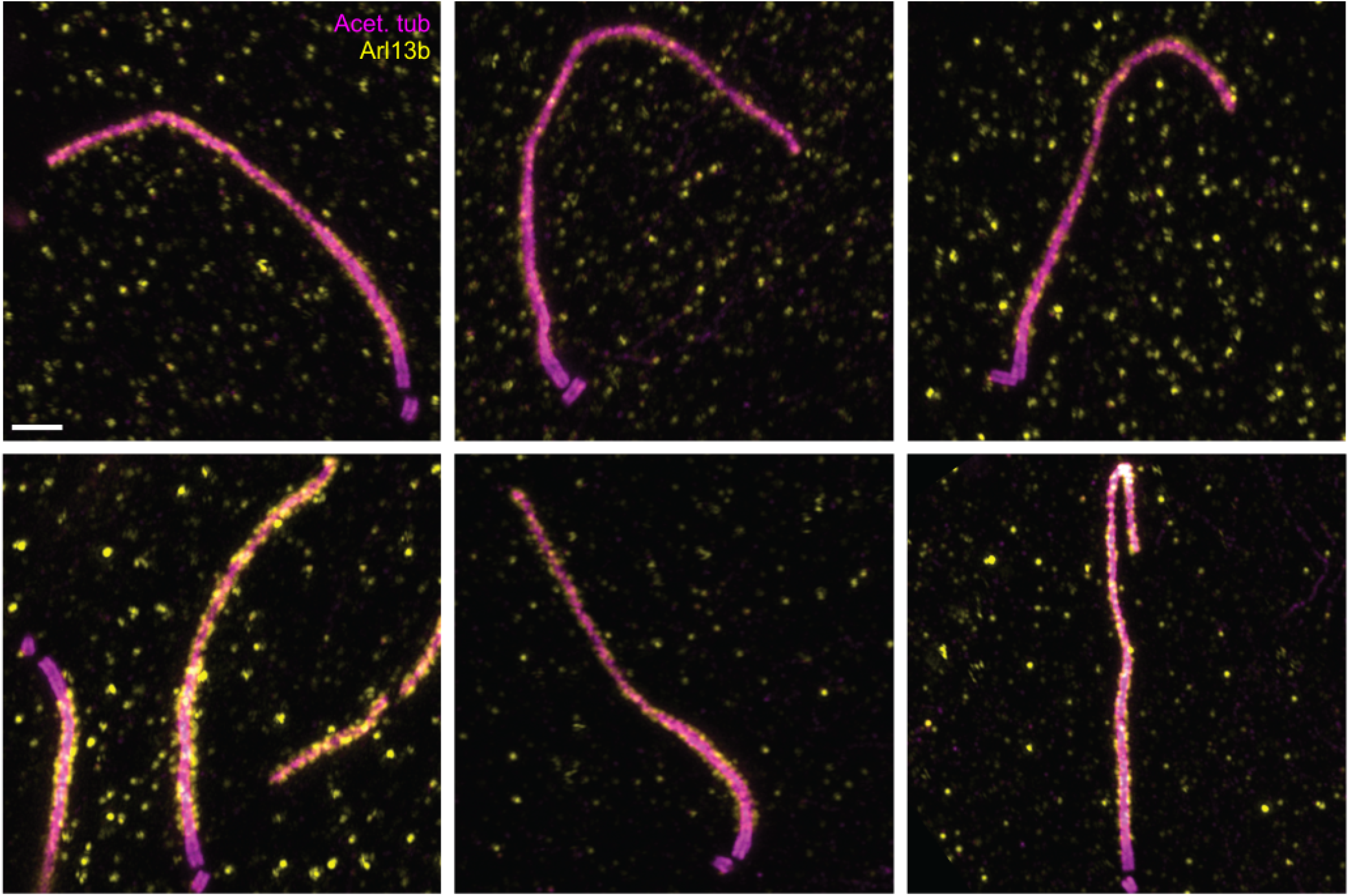
Variability in islet cell primary cilia curvature in UExM images. Broken islet with visible edges and microvilli of the islet cells as well as primary cilia (purple). Scale bars: 1 μm.

**Supplementary Figure 4.**
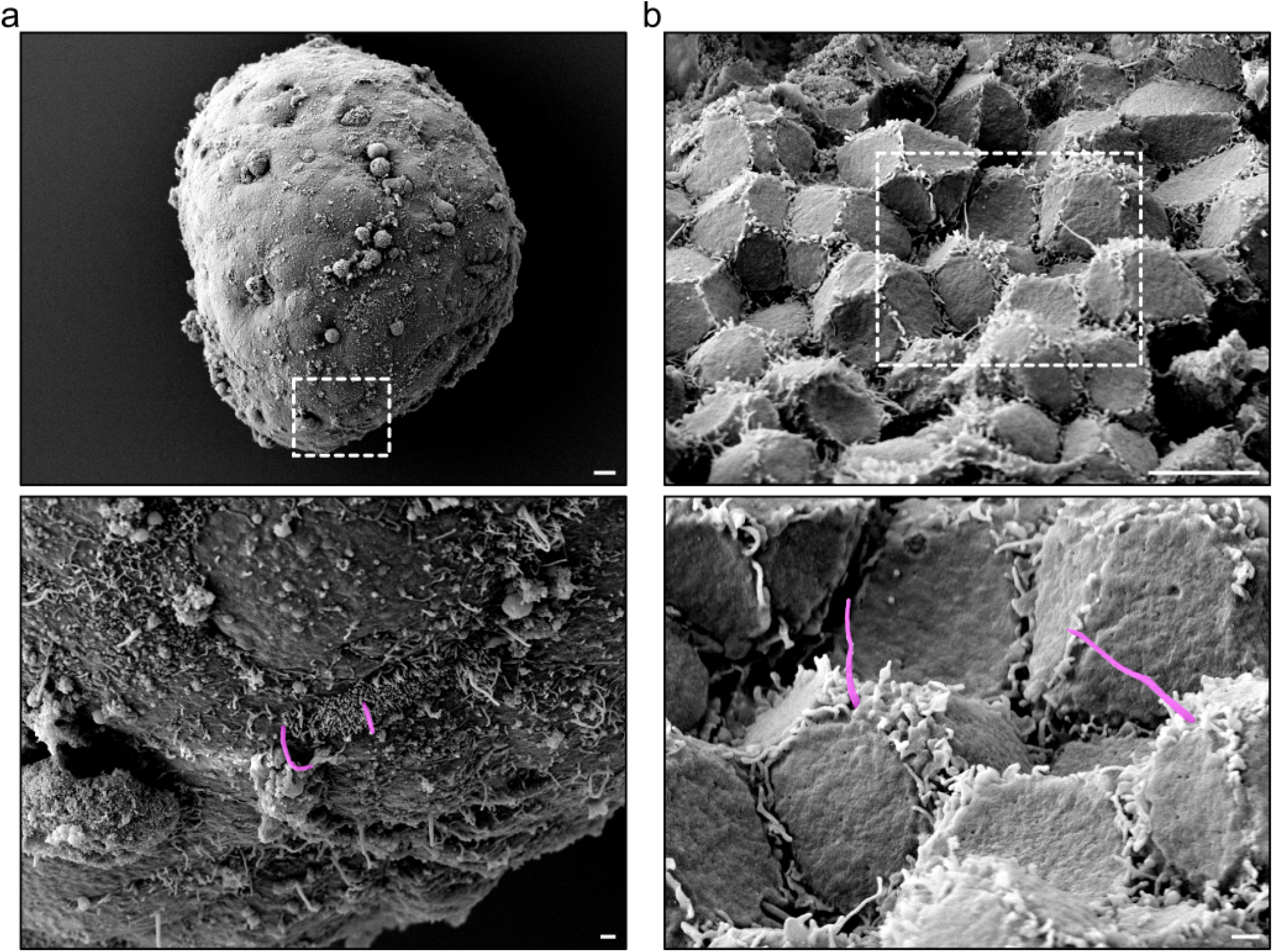
SEM of isolated intact and broken mouse islets. **a)** Intact islet with magnification of boxed area showing islet cell primary cilia (purple). **b)** Broken islet with visible edges and microvilli of the islet cells as well as primary cilia (purple). Scale bars: 1 μm.

**Supplementary Figure 5.**
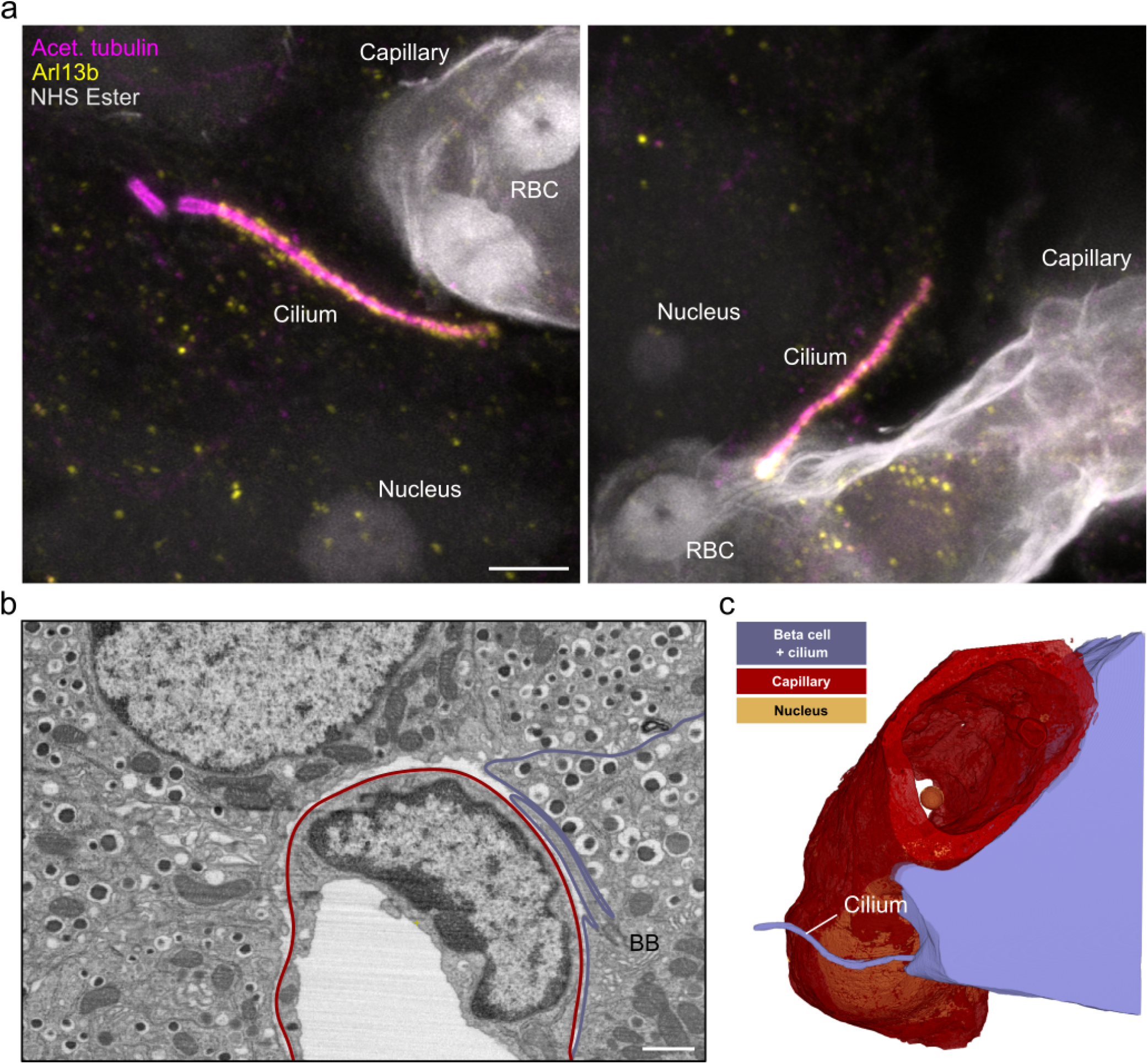
Interaction of islet cell primary cilia with blood vessels. **a)** Expanded beta cell cilia in close contact with blood vessels, as visualized by acetylated tubulin (magenta) Arl13b (yellow) and NHS Ester (gray). Scale bar: 1 μm. **b)** FIB-SEM slice showing a blood vessel with adjacent islet cells. A beta cell projects its primary cilium along the endothelial cells forming the vessel. Scale bar: 500 nm. **c)** The 3D rendering shows the primary cilium projecting along the vessel.

**Supplementary Figure 6.**
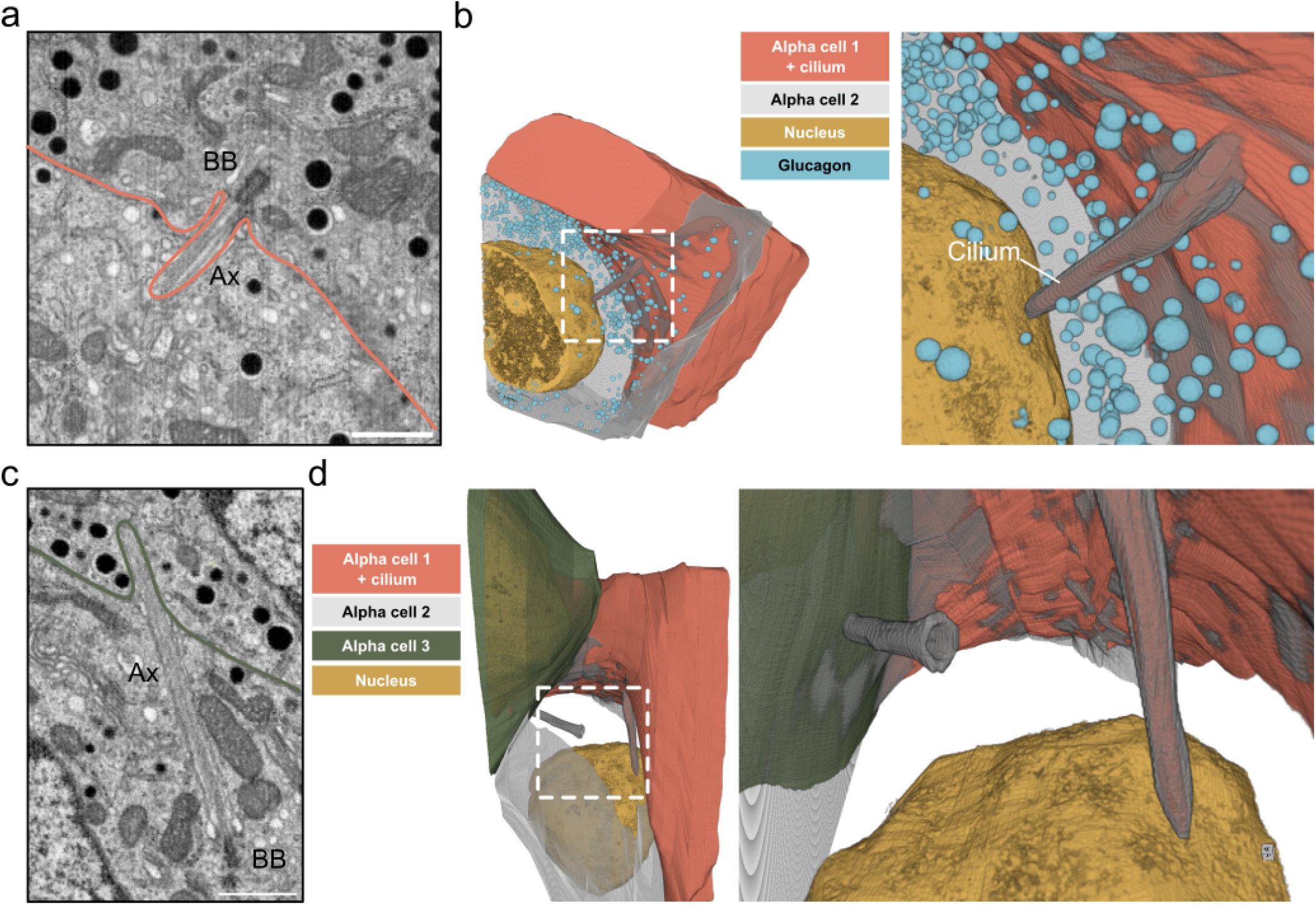
Alpha cell primary cilium “pinches” a neighboring alpha cell. **a)** FIB-SEM slice showing an alpha cell primary cilium with basal body (BB) and axoneme) pushing into a neighboring alpha cell. Scale bar: 500 nm **b)** 3D rendering of both cells with inset depicting the termination of the primary cilium close to the nucleus. **c)** A FIB-SEM slice through the gray alpha cell depicted in b) with its cilium pinching into the neighboring alpha cell (green). Scale bar: 500 nm. **d)** 3D renderings of all 3 alpha cells with 2 cilia protruding into the adjacent cell.

**Supplementary Figure 7.**
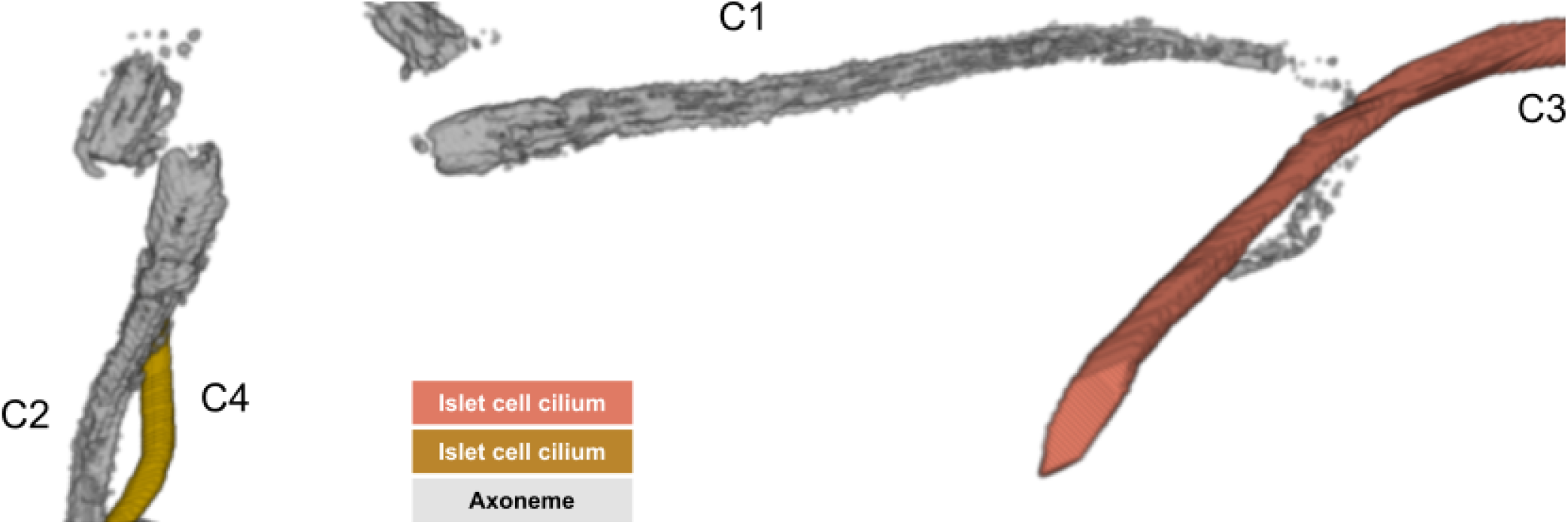
Cilia in contact with neighboring cilia. 3D renderings of the two primary cilia (C1 and C2) of a double-ciliated beta cell which are in close proximity to cilia (C3 and C4) from neighboring islet cells on opposite sides of the beta cell. Smaller images show magnified renderings with the plasma membrane of the beta cell.

**Supplementary Figure 8.**
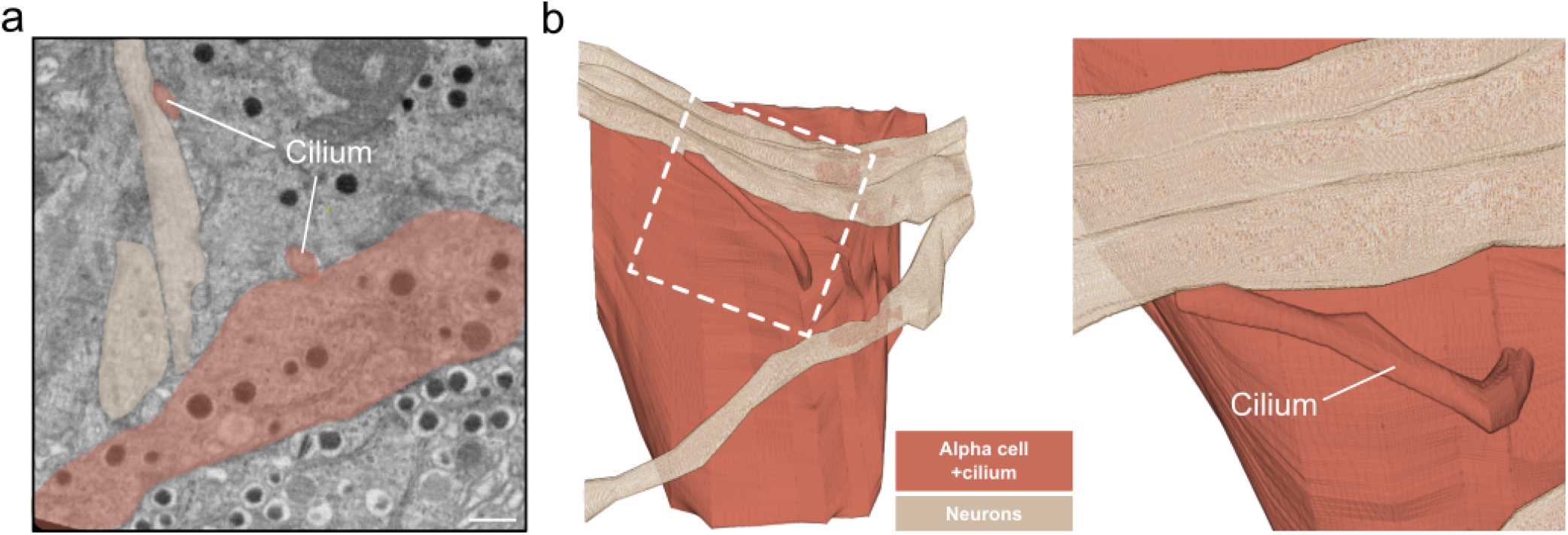
Connection of alpha cell primary cilium tip with neurons. **a)** Slice through a FIB-SEM volume with nerves (beige) and an alpha cell and its cilium (orange). Scale bar: 500 nm **b)** 3D renderings of nerves and the alpha cell. The magnified view shows the connections between the cilium tip and the neuron.

## Supplementary Note 2: Videos

**Video abstract**.

**Video 1: Raw ssET of a human beta cell primary cilium**.

**Video 2: FIB-SEM + Axoneme reconstruction of a mouse beta cell primary cilium. Video 3: Spatial restriction of beta cell primary cilia**.

**Video 4: Beta cell cilium synapse**.

